# The endoplasmic reticulum autophagy receptor TEX264 drives epidermal differentiation and is dysregulated in Darier disease

**DOI:** 10.1101/2025.08.05.668774

**Authors:** Christopher J. Johnson, Afua Tiwaa, Arti Parihar, Edwin Antonio, Brooke D. Lorenz, Reeteka Kudallur, Aaron Ramonett, Anthony Coon, Mrinal K. Sarkar, Johann E. Gudjonsson, Cory L. Simpson

**Affiliations:** Department of Dermatology, University of Washington, Seattle, WA, USA; University of Washington School of Medicine, Seattle, WA, USA; Department of Laboratory Medicine & Pathology, University of Washington, Seattle, WA, USA; Department of Dermatology, University of Michigan, Ann Arbor, MI, USA; Institute for Stem Cell & Regenerative Medicine, University of Washington, Seattle, WA, USA

## Abstract

Differentiating keratinocytes break down their organelles and nuclei to become the compacted cornified layers of the epidermal barrier in a poorly understood catabolic process. Live confocal imaging of stratified human organotypic epidermis revealed endoplasmic reticulum (ER) fragmentation and lysosomal engulfment in the cornifying layers, where we found up-regulation of TEX264, a receptor that mediates selective autophagy of the ER (reticulophagy). TEX264 expression was increased by ER stress, which caused precocious cornification of organotypic epidermis. In undifferentiated keratinocytes, ectopic TEX264 was sufficient to fragment the ER, while in highly differentiated keratinocytes, it accelerated ER elimination and induced nuclear shrinkage; these effects were abolished by mutating the LC3 interacting region required for its autophagic function. Knockout of TEX264 or inhibiting its activation disrupted maturation of organotypic cultures, pointing to a critical role for reticulophagy in cornification. Finally, in patient biopsies and an organotypic model of Darier disease, a genetic cornification disorder linked to ER dysfunction, we found increased TEX264 in areas of premature cornification (dyskeratosis). Our results identified TEX264 as a key driver of epidermal differentiation and led us to propose a novel model of cornification in which keratinocytes activate selective autophagy receptors to orchestrate orderly organelle elimination during cutaneous barrier formation.

**GRAPHICAL ABSTRACT:** 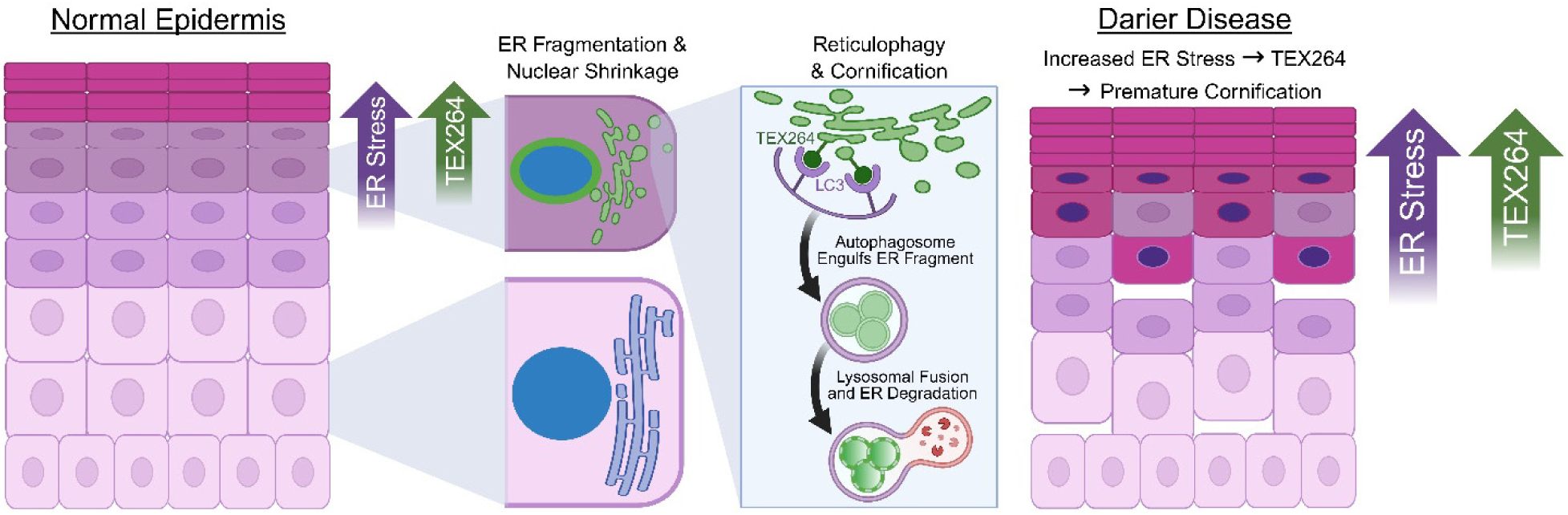

## INTRODUCTION

The epidermis constitutes the outermost portion of the skin and defends the body against microbes, wounds, and water loss (1). To generate the epidermal barrier, multi-layered keratinocytes undergo a unique terminal differentiation program as they move outward in the epithelium (2, 3). In a specialized catabolic process termed cornification, they ultimately transition into corneocytes upon eliminating their organelles and nuclei, which allows them to become highly compressed in the superficial dead skin layers (4–6). The critical nature of this process is underscored by numerous genetic disorders of cornification (7–9), most of which lack any targeted therapy since the mechanisms governing organelle and nuclear breakdown during keratinocyte maturation remain poorly understood (10, 11).

While morphological changes in mitochondria during epidermal differentiation have been more straightforward to visualize in both classical electron microscopy (12–14) and live imaging studies (15, 16), the ultimate fate of the endoplasmic reticulum (ER) in cornifying keratinocytes remained unknown. Since ER is the primary intracellular store of calcium, a major driver of epidermal differentiation (17–19), and was shown to intimately associate with intercellular junctions (20), dismantling the ER tubular network during keratinocyte maturation must be a highly regulated process. Underscoring the importance of this organelle in human epidermis, dysfunction of an ER calcium pump has been linked to a genetic disorder of cornification called Darier disease (21–23). Our previous work established that mitochondria undergo fragmentation and routing into lysosomes during cornification via the selective autophagy pathway utilizing a mitochondria-tethered receptor NIX (16). This discovery prompted us to ask if a parallel pathway is required to selectively degrade the ER during the final stage of keratinocyte maturation.

Recent work identified several transmembrane protein receptors that drive selective autophagy of the ER (termed reticulophagy or ER-phagy) in various cell types (24, 25), though they had not been studied in the epidermis. The best-characterized reticulophagy receptors include CCPG1, FAM134B, RTN3, SEC62, and TEX264, which are embedded in the outer ER surface and can induce membrane curvature and organelle fragmentation to enable their degradation (26, 27). Each receptor contains a conserved LC3-interacting region (LIR) that binds the LC3 ligand on autophagosomal membranes, which encircle the organelle cargo (28). Double-membrane autophagosomes ultimately fuse with lysosomes to acidify and degrade their contents (29). We hypothesized that the ER in differentiating keratinocytes is degraded through selective reticulophagy in the outermost epidermal layers to form the cornified skin barrier.

## RESULTS

### Keratinocyte ER undergoes progressive fragmentation and lysosomal acidification during epidermal differentiation

To replicate the process of epidermal stratification and differentiation in a tractable 3D *in vitro* platform compatible with live imaging, we utilized an organotypic tissue model that generates mature epidermis from human keratinocytes grown at an air-liquid interface (30).

Upon reaching the outermost layers of the epidermal tissue, the keratinocytes eliminate membrane-bound organelles and nuclei to form a mature stratum corneum after 5-7 days. We optimized the growth and differentiation of keratinocytes in both submerged and organotypic cultures for live imaging of fluorophore-tagged organelles by spinning-disk confocal (SDC) microscopy (16), allowing us to illuminate the mechanisms of skin barrier formation.

To track the remodeling and fate of the ER, a highly dynamic organelle (31), in a 3D epidermal tissue, we used live normal human epidermal keratinocytes (NHEKs) or TERT-immortalized human epidermal keratinocytes (THEKs) transduced with pH-stable fluorophores targeted to the ER lumen (ER-dsRed; ER-StayGold (32)) or the ER outer membrane (mChilada-VAPB). We then obtained Z-stack images through the entire organotypic epidermis grown from stably transduced keratinocytes (**Figure 1A**). While the ER in the lower layers showed a typical tubular network, as keratinocytes differentiated and moved upward in the epithelium, we noted the appearance of ER puncta, indicating organelle breakdown (**Figure 1B**). Remarkably, in the outermost epidermal layers undergoing cornification, the ER network completely fragmented.

**Figure 1:**
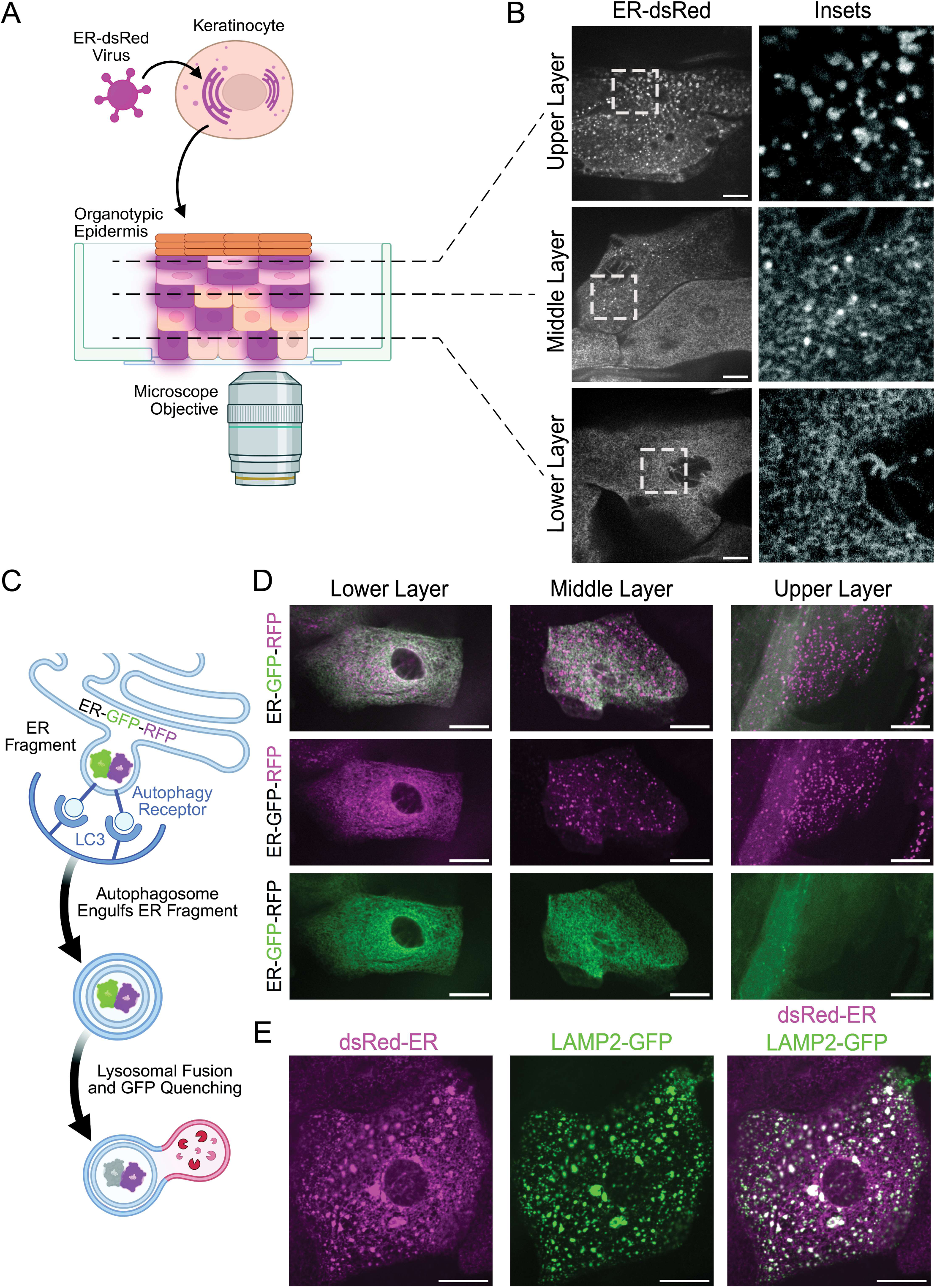
Keratinocyte ER undergoes progressive fragmentation and lysosomal acidification during epidermal differentiation. (**A**) Diagram depicting spinning-disk confocal (SDC) microscopy of live organotypic epidermis grown from keratinocytes transduced with a fluorophore (dsRed) targeted to the ER. (**B**) Live SDC microscopy images from the lower, middle, and upper layers of ER-dsRed-transduced organotypic epidermis grown from NHEKs displaying progressive breakdown of the tubular ER network with accumulation of circular ER fragments; bar = 10 µm; insets magnified at right. (**C**) Diagram of ER-GFP-RFP labeling an ER fragment that is engulfed by LC3-labeled autophagosomal membrane, then has its GFP signal quenched by acidic pH due to autophagosome-lysosome fusion, while the RFP signal persists. (**D**) Live SDC microscopy images from the lower, middle, and upper layers of organotypic epidermis grown from NHEKs transduced with ER-GFP-RFP reveal progressive loss of GFP in the upper layers with ER fragments having only RFP signal; bar = 10 µm. (**E**) Live SDC microscopy images of the upper layers of organotypic epidermis grown from NHEKs transduced with both ER-dsRed and LAMP2-GFP, demonstrating localization of ER fragments within lysosomes; bar = 10 µm.

We next sought to determine if ER fragments are routed into acidic lysosomes using organotypic rafts grown from NHEKs transduced with ER-GFP-RFP, a tandem fluorophore targeted to the ER lumen that acts as a pH sensor: in a neutral environment, both GFP and RFP fluoresce, but when routed into a low pH environment, the GFP signal is quenched (**Figure 1C**). SDC images demonstrated both red and green signals in the tubular ER network within the lower keratinocyte layers, while GFP-negative ER fragments began to appear in the middle layers (**Figure 1D**). The upper epidermal layers demonstrated loss of GFP signal in the ER along with complete remodeling of ER tubules into spherical ER fragments positive for RFP only, implying the organelle remnants were routed into an acidic compartment.

To confirm these GFP-negative ER puncta were localized in lysosomes, we grew organotypic epidermis from NHEKs co-transduced with GFP tethered to the outer lysosomal surface (LAMP2-GFP) along with ER-dsRed. Live confocal imaging of these cultures allowed us to demonstrate that dsRed-tagged ER fragments in the upper epidermal layers were contained within GFP-labeled lysosomes (**Figure 1E**). These data indicate that the ER undergoes progressive breakdown during keratinocyte differentiation, generating organelle fragments that are targeted for lysosomal degradation. This led us to hypothesize that keratinocytes route their ER into lysosomes using the autophagy machinery. We next sought to identify the mechanism that allowed keratinocytes to target the ER for autophagic degradation in a selective and differentiation-dependent manner.

### The autophagy receptor TEX264 is upregulated during keratinocyte differentiation and fragments the ER in an LC3-dependent manner

As opposed to macro-autophagy, which drives nonselective degradation of cytoplasmic contents (33), selective autophagy specifically removes damaged or superfluous organelles (34, 35). Cells utilize selective autophagy of the ER for organelle remodeling and quality control, degrading dysfunctional portions (24, 27, 28). ER damage induces expression of reticulophagy receptors on the outer ER membrane; these receptors bind to Atg8-related proteins like LC3 on precursor autophagosomal membranes, which engulf fragments of the ER and route them into lysosomes for breakdown (26). To define whether and where reticulophagy receptors are expressed in the skin, we queried a public single-cell murine skin RNA sequencing (RNAseq) database (https://linnarssonlab.org/epidermis/) to identify a candidate receptor that could drive ER breakdown during keratinocyte differentiation (36). Intriguingly, we found TEX264, a reticulophagy receptor (37, 38), was markedly induced in the uppermost viable layers of the epidermis, which are preparing to undergo cornification (**Figure 2A**).

**Figure 2:**
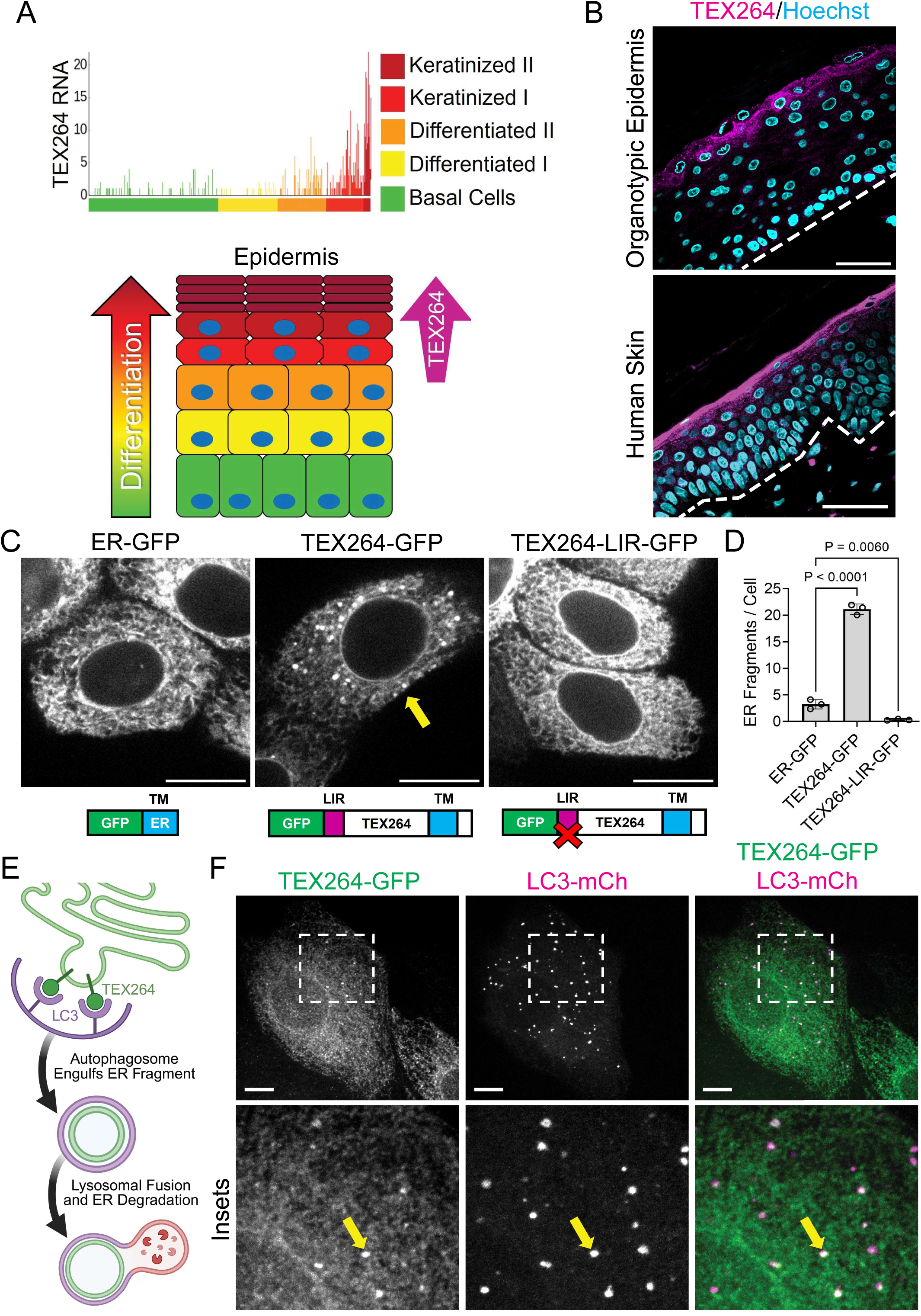
The autophagy receptor TEX264 is upregulated during keratinocyte differentiation and fragments the ER in an LC3-dependent manner. (**A**) Graph of TEX264 mRNA expression from the public single-cell RNA sequencing database from mouse epidermal keratinocytes (http://linnarssonlab.org/epidermis/) (36); (below) color-coded diagram depicting keratinocytes progressively differentiating within the epidermis with a marked increase in TEX264 mRNA in the uppermost layers. (**B**) Immunostaining of TEX264 (magenta) and nuclei (Hoechst; cyan) in tissue cross-sections from organotypic epidermis grown from NHEKs (representative of N=5 unrelated donors) and a patient skin biopsy (representative of N=10 unrelated donors); bar = 50 µm; dashed lines mark bottom of the epidermis. (**C**) SDC fluorescence microscopy images of NHEKs transduced with ER-GFP, TEX264-GFP or TEX264-LIR-GFP (bar = 10 µm); yellow arrow points to an ER fragment positive for TEX264-GFP; (below) diagrams of each ER-targeted protein construct including the GFP tag (green), LC3 interacting region (LIR, magenta), and transmembrane domain (TM, blue). (**D**) Bar graph displays the mean ± SD of the number of ER fragments per cell (for ≥70 cells per condition) from N=3 biological replicates; *P* values are from 1-way ANOVA using the Dunnett adjustment for multiple comparisons. (**E**) Diagram of TEX264-GFP on the surface of an ER fragment; TEX264 binding to LC3 on the autophagosomal membrane engulfs the ER fragment that then fuses with a lysosome to degrade the ER remnant. (**F**) SDC microscopy images of NHEKs co-transduced with TEX264-GFP and LC3-mCherry (LC3-mCh); bar = 10 µm; insets magnified below; yellow arrows highlight TEX264-GFP and LC3-mCh colocalized in ER fragments.

Prior to our study, TEX264 had not been investigated in the skin; thus, we first confirmed the localization of endogenous TEX264 protein by immuno-staining both organotypic epidermis and human skin biopsies. Consistent with murine skin RNAseq data, TEX264 was concentrated in the upper (granular) keratinocyte layers (**Figure 2B**), coinciding with the location of fragmentation and lysosomal routing of the ER that we observed in live organotypic epidermis. Building on this circumstantial evidence, we tested if TEX264 played an active role in ER breakdown by transducing undifferentiated NHEKs (lacking endogenous TEX264) with a GFP-tagged receptor. Ectopic TEX264-GFP was sufficient to induce fragmentation of the ER network (**Figure 2C-D**) compared to GFP alone targeted to the ER (ER-GFP).

To further rule out non-specific effects on ER morphology, we transduced NHEKs with TEX264 having its LIR mutated (TEX264-LIR-GFP) to prevent LC3 binding (37, 39). This completely dampened ER fragmentation, demonstrating that the ability to engage the autophagy machinery is essential to the function of TEX264 in ER breakdown (**Figure 2E**). To confirm that TEX264-positive ER remnants were routed into autophagosomes, we co-transduced NHEKs with TEX264-GFP and the mCherry-tagged autophagosomal marker LC3B (LC3-mCh), which colocalized with the ER fragments (**Figure 2F**). Together, these observations confirm that TEX264 functions as a reticulophagy receptor in epidermal keratinocytes. We next sought to understand what drives the spatiotemporal control of TEX264 expression in the epidermis to appropriately trigger the process of ER breakdown and cornification.

### ER stress upregulates TEX264 in organotypic epidermis and induces premature terminal differentiation

ER stress is a potent stimulator of reticulophagy in many cell types in order to degrade damaged portions of the organelle or misfolded proteins (24, 40). Interestingly, it has been shown that ER stress levels rise in the upper layers of the epidermis and promote keratinocyte differentiation (41, 42). This known timing of epidermal ER stress correlated strongly with our immunostaining localizing TEX264 expression to the granular layers of the epidermis (**Figure 2B**). Thus, we proposed that ER stress upregulates TEX264 to degrade the ER and trigger cornification in the uppermost keratinocyte layers. We tested this potential relationship between ER stress and TEX264 by treating organotypic epidermis with tunicamycin for 48 hr.

Sustained ER stress from tunicamycin caused premature cornification, which reduced epidermal tissue thickness due to acceleration of the cornification process (**Figure 3A-B**). This phenotype was paralleled by precocious TEX264 expression in the immediate supra-basal layers (**Figure 3C-D**). The treated keratinocytes, which normally form several intermediate (spinous) layers, instead abruptly transitioned into a granular layer phenotype marked by basophilic keratohyalin granules in H&E-stained tissue sections and underwent premature cornification. These data suggest that ER stress accumulation in the upper epidermal layers augments TEX264 levels and led us to test whether TEX264 is sufficient to drive cornification.

**Figure 3:**
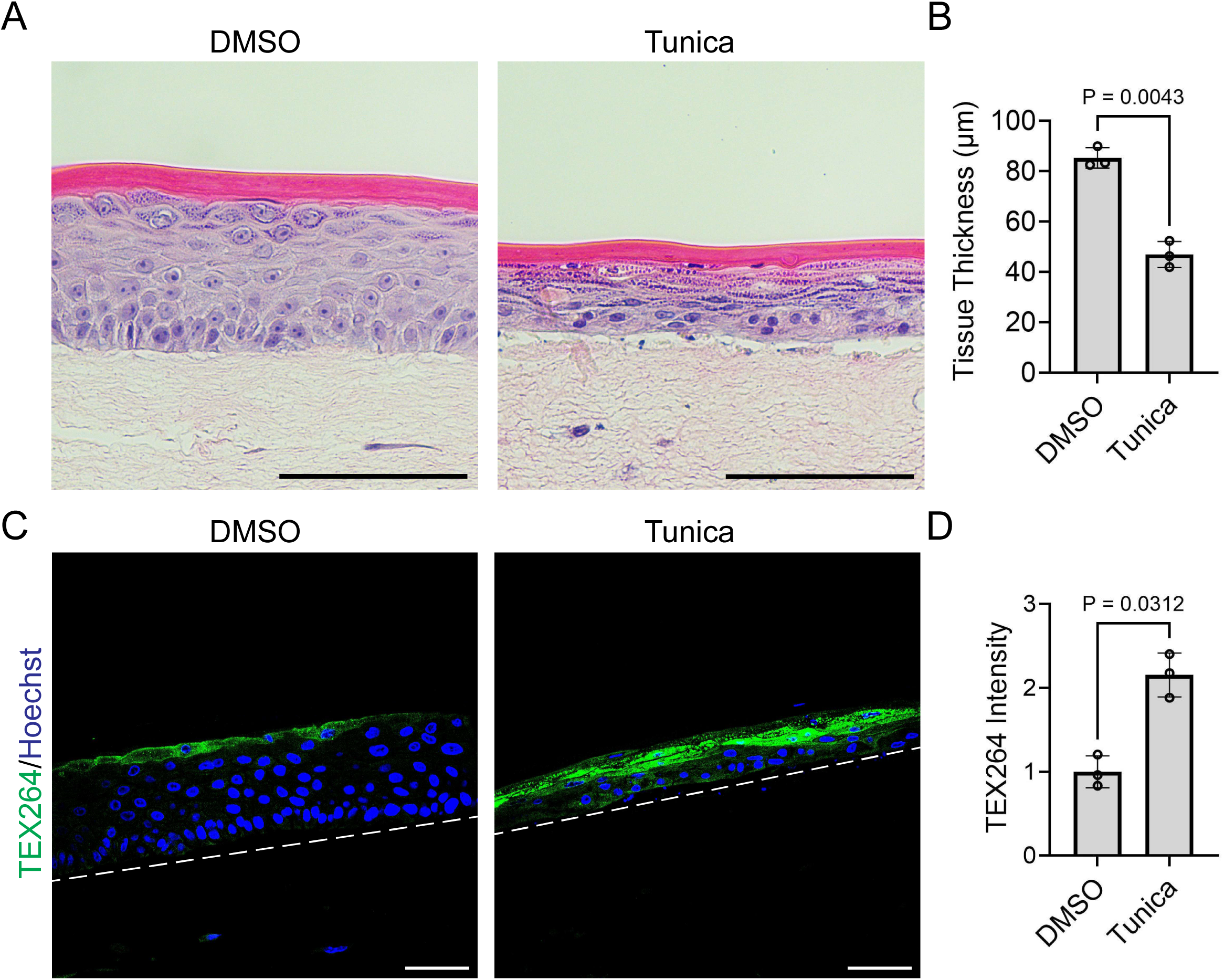
ER stress upregulates TEX264 in organotypic epidermis and induces premature terminal differentiation. (**A**) H&E-stained cross-sections of organotypic epidermal cultures treated with DMSO or tunicamycin (Tunica) for 48 hr; bar = 100 µm. (**B**) Bar graph displays the mean ± SD of the tissue thickness (µm) of epidermal cultures treated with DMSO or Tunica with individual data points representing the average thickness across N=3 biological replicates; *P* values from a ratio paired 2-tailed Student’s t test. (**C**) Immunostaining of TEX264 (green) and nuclei (Hoechst, blue) in cross-sections of organotypic epidermal cultures treated with DMSO or Tunica for 48 hr; bar = 50 µm; dashed lines mark bottom of the epidermis. (**D**) Bar graph displays the mean ± SD of TEX264 fluorescence intensity in cultures treated with DMSO or Tunica with data points from N=3 biological replicates; mean intensity for DMSO normalized to 1; *P* value from a paired 2-tailed Student’s t test.

### Ectopic TEX264 accelerates ER breakdown and induces nuclear shrinkage in differentiating keratinocytes

To elucidate the function of TEX264 during later stages of keratinocyte maturation, we differentiated keratinocytes transduced with TEX264-GFP. We grew these cells to confluency in low-calcium (0.3 mM) serum-free medium, then switched them into 3D differentiation medium. After 4 days, cells were fixed and immunostained. To track ER fate in keratinocytes expressing ER-GFP or TEX264-GFP, we immunostained calreticulin, a chaperone protein found in the ER lumen. Keratinocytes over-expressing TEX264-GFP exhibited increased ER degradation with a marked loss of calreticulin compared to ER-GFP (**Figure 4A-B**), confirming the reticulophagy receptor is sufficient to promote ER breakdown in highly differentiated keratinocytes.

**Figure 4:**
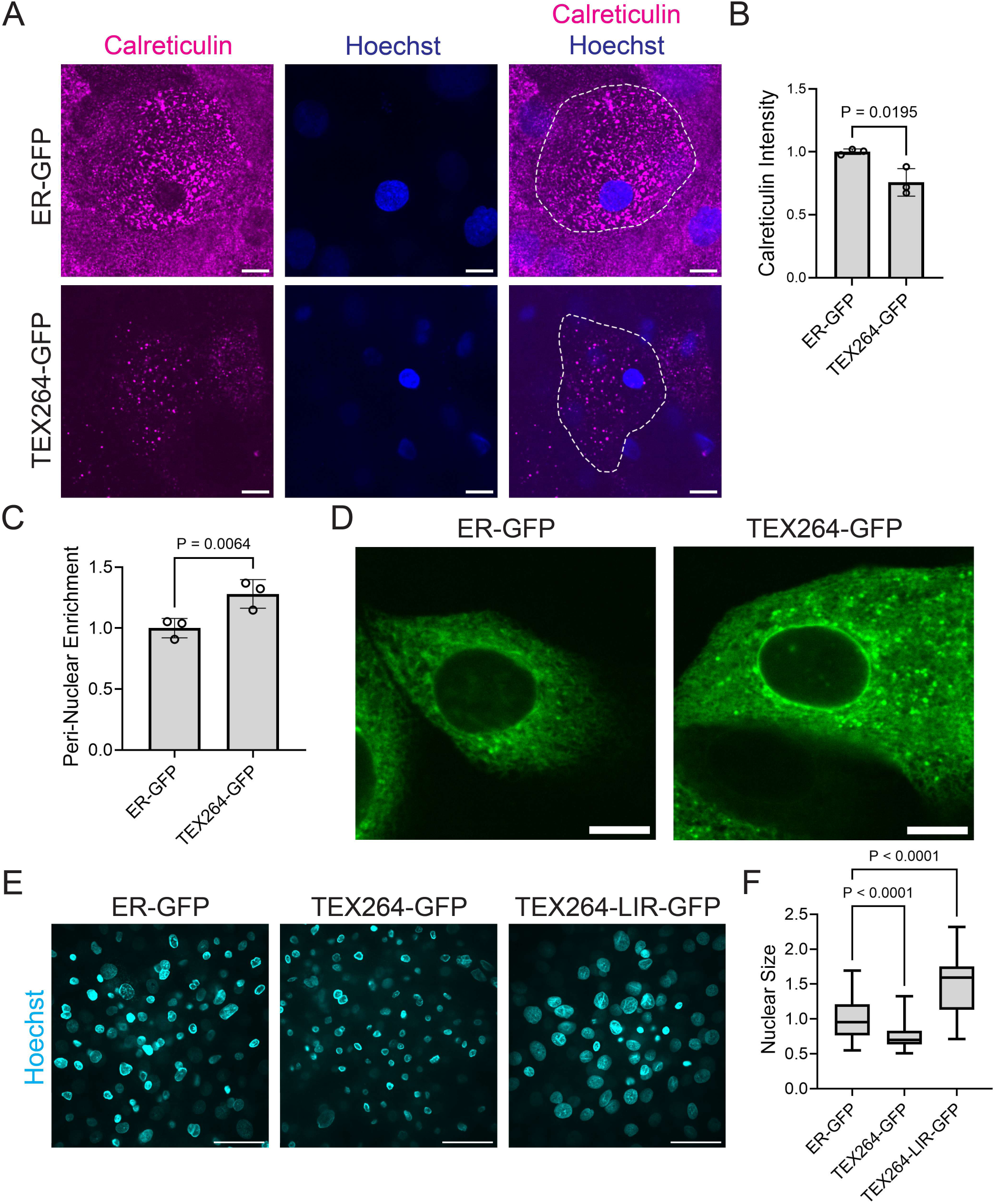
Ectopic TEX264 accelerates ER breakdown and induces nuclear shrinkage in differentiating keratinocytes. (**A**) SDC microscopy images of THEKs transduced with either ER-GFP or TEX264-GFP that were differentiated in 3D medium for 4 days, then stained with the ER marker calreticulin (magenta) and Hoechst (blue); dashed lines surround a single cell; bar = 10 µm. (**B**) Bar graph displays the mean ± SD of calreticulin intensity per cell (from ≥23 images per condition) with individual data points from N=3 biological replicates; mean intensity for ER-GFP was normalized to 1; *P* values are from an unpaired 2-tailed Student’s t test. (**C**) Bar graph displays the mean ± SD of the perinuclear enrichment factor (ratio of perinuclear to cytoplasmic GFP intensity) of NHEKs transduced with either ER-GFP or TEX264-GFP with individual data points representing N=3 biological replicates; mean perinuclear enrichment for ER-GFP was normalized to 1; *P* values are from a paired 2-tailed Student’s t test. (**D**) SDC microscopy images of NHEKs transduced with either ER-GFP or TEX264-GFP highlighting the perinuclear enrichment of TEX264-GFP; bar = 10 µm. (**E**) SDC microscopy images of THEKs transduced with ER-GFP, TEX264-GFP, or TEX264-LIR-GFP that were differentiated in 3D medium for 4 days, then stained with Hoechst (cyan); bar = 50 µm. (**F**) Nuclear sizes are shown as a box plot of the 25^th^-75^th^ percentile with a line at the median from N≥60 images per condition from 3 biological replicates of THEKs transduced with ER-GFP, TEX264-GFP, or TEX264-LIR-GFP; mean nuclear size for ER-GFP was normalized to 1; *P* values are from 1-way ANOVA using the Dunnett adjustment for multiple comparisons.

We also observed perinuclear enrichment of TEX264-GFP in keratinocytes (**Figure 4C-D**), consistent with studies in other cell types (43, 44). Its transmembrane targeting sequence localizes the receptor to the outer ER membrane, which is continuous with the outer leaflet of the nuclear envelope. Intriguingly, we noted that highly differentiated keratinocytes expressing ectopic TEX264 exhibited reduced nuclear size. When quantified, we found TEX264-GFP induced shrinkage of nuclei (**Figure 4E-F**), which occurs in the uppermost epidermal layers as they prepare to cornify. This function of TEX264 depended on its ability to engage the autophagy machinery since cells expressing TEX264-LIR-GFP actually exhibited an increase rather than a reduction in nuclear size. These findings led us to propose that TEX264 participates in nuclear degradation during the terminal stage of keratinocyte differentiation.

### Halting autophagic degradation of TEX264-positive ER fragments disrupts epidermal differentiation

We next tested if blocking TEX264 function disrupted epidermal differentiation. Previous work identified the LIR domain of TEX264 that is crucial for binding LC3 (37), but investigators also found critical serine residues upstream of the LIR domain. These are specifically phosphorylated by casein kinase 2 (CK2) to prime the reticulophagy receptor for activation by enhancing its affinity for LC3 (39). To inhibit TEX264 activation, we treated keratinocytes transduced with TEX264-GFP with the CK2 inhibitor CX-4945 (CX). Compared to DMSO, we found CK2 inhibition increased the number of TEX264-positive ER fragments, suggesting they cannot be efficiently routed for autophagic degradation (**Figure 5A-B**). We noted similar accumulation of TEX264-positive ER fragments upon blockade of autophagosome-lysosome fusion and acidification with bafilomycin A1 (BafA1). In contrast, similar effects on ER fragments were not seen in cells expressing the LC3-uncoupled mutant of TEX264 (TEX264-LIR-GFP), which lacks an activatable LIR domain.

**Figure 5:**
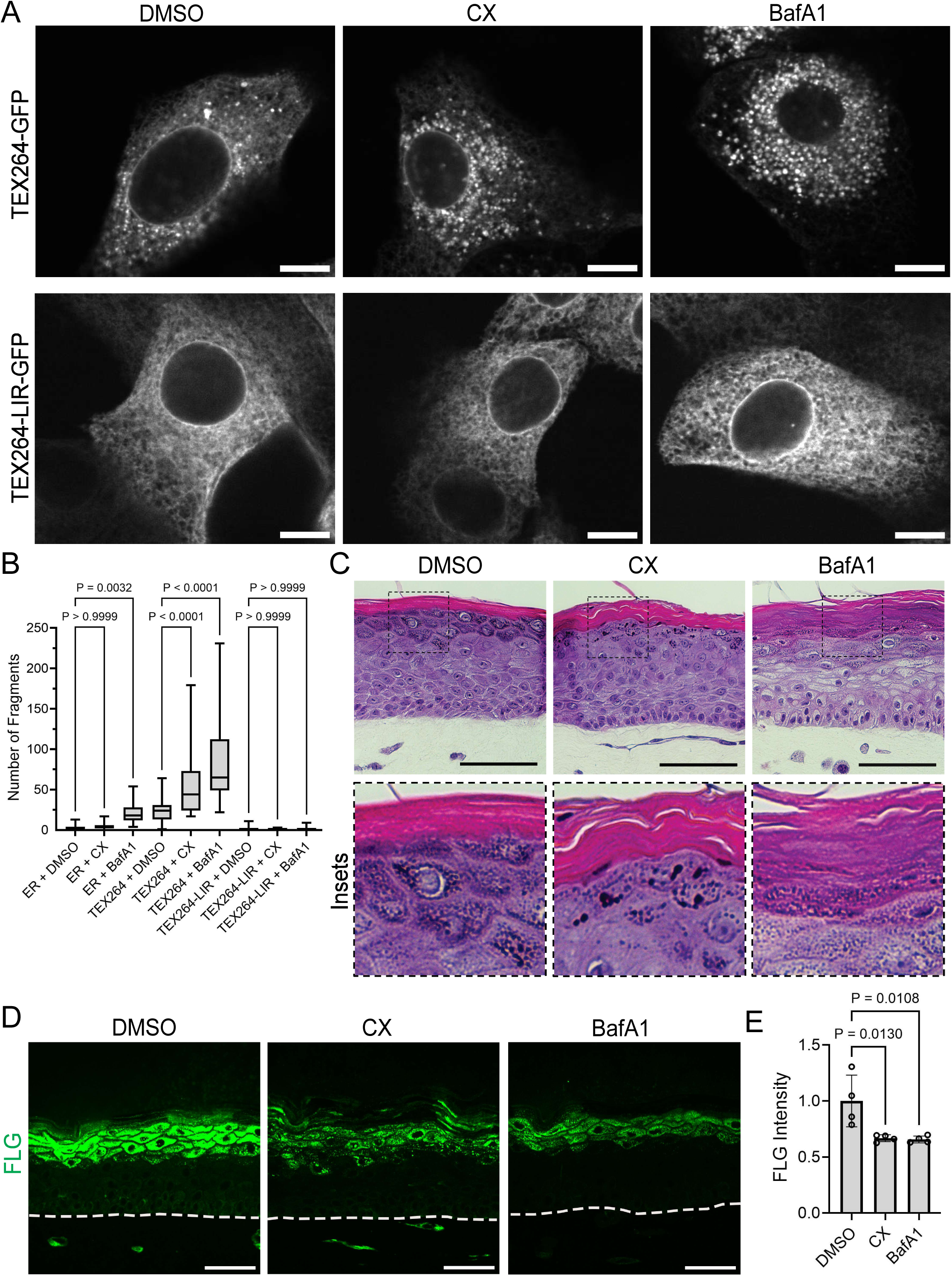
Halting autophagic degradation of TEX264-positive ER fragments disrupts epidermal differentiation. (**A**) SDC microscopy images showing ER fragments in live undifferentiated NHEKs transduced with TEX264-GFP or TEX264-LIR-GFP and treated with DMSO, CX-4945 (CX), or bafilomycin-A1 (BafA1); bar = 10 µm. (**B**) Number of ER fragments per cell are shown as a box plot of the 25^th^-75^th^ percentile with a line at the median from N≥38 images per condition from 3 biological replicates of NHEKs transduced with ER-GFP, TEX264-GFP or TEX264-LIR-GFP and treated with DMSO, CX, or BafA1; *P* values are from 1-way ANOVA using the Bonferroni adjustment for multiple comparisons. (**C**) H&E-stained cross-sections of organotypic epidermal cultures of NHEKs treated with DMSO, CX, or BafA1 for 48 hr (insets magnified below); bar = 100 µm. (**D**) Images of immunostaining for filaggrin (FLG, green) in cross-sections of organotypic epidermal cultures treated with DMSO, CX, or BafA1; bar = 50 µm; dashed lines mark bottom of the epidermis. (**E**) Bar graph displays the mean ± SD of FLG intensity of organotypic epidermal cultures treated with DMSO, CX, or BafA1 with individual data points from N=4 biological replicates; mean FLG intensity for DMSO was normalized to 1; *P* values are from 1-way ANOVA using the Dunnett adjustment for multiple comparisons.

Given our hypothesis that TEX264 serves as a critical driver of keratinocyte differentiation, we tested whether inhibiting activation or degradation of endogenous TEX264 using CX or BafA1, respectively, would impair epidermal morphogenesis. Treatment of organotypic epidermis with CX resulted in abnormal keratinocyte maturation with abnormal formation of keratohyalin granules and nuclear retention in the cornified layers (**Figure 5C, insets**). A more dramatic effect was seen with BafA1, which non-selectively blocks all autophagic flux. This drug grossly impaired epidermal maturation with vacuole accumulation in lower layers, fewer and smaller keratohyalin granules, and an abnormal transition of cells into the cornified layers, which exhibited impaired compaction and retention of nuclei. Given these phenotypes, we performed immunostaining in CX- and BafA1-treated epidermal cultures for filaggrin (FLG), a key marker of the granular layer that aids in keratinocyte compaction (45–47). Compared to DMSO-treated controls, we found a marked reduction in FLG levels in both CX- and BafA1-treated organotypic epidermis (**Figure 5D-E**), confirming that blocking CK2, which is required to activate TEX264, impaired keratinocyte maturation.

### TEX264 knockout impairs maturation of keratinocytes within the upper epidermal layers

To more specifically test whether TEX264 is necessary for keratinocyte differentiation, we utilized CRISPR/Cas9 to engineer multiple TEX264 knockout (KO) lines of TERT-immortalized human epidermal keratinocytes (THEKs). We used two independent TEX264-deficient cell lines (KO-1 and KO-2) to grow organotypic epidermal cultures (**Figure 6A**); these were compared to control cells having a pseudogene similarly targeted by CRISPR/Cas9.

**Figure 6:**
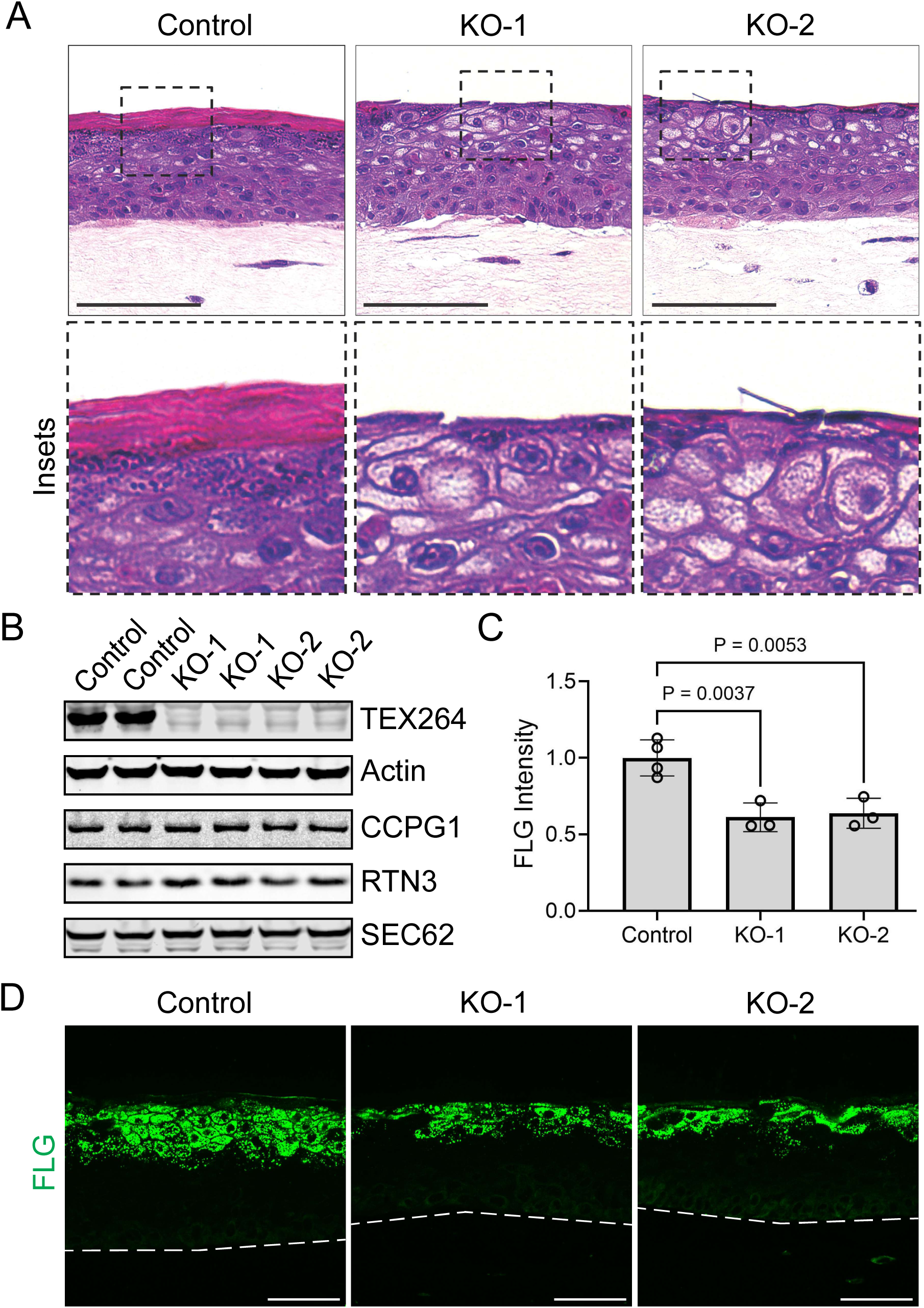
TEX264 knockout impairs maturation of keratinocytes within the upper epidermal layers. (**A**) H&E-stained cross sections of organotypic epidermal cultures grown from control or 2 independent lines of TEX264 KO THEKs; KO-1 and KO-2 cultures displayed impaired granular layer maturation and cornification (insets magnified below); bar = 100 µm. (**B**) Immunoblotting of control and KO THEK lysates for the reticulophagy receptors TEX264, CCPG1, RTN3, and SEC62; actin serves as a loading control. (**C**) Bar graph displays the mean ± SD of FLG intensity of organotypic epidermal cultures grown from control or KO THEKs with individual data points from N=3 or 4 biological replicates; mean FLG intensity for control cultures was normalized to 1; *P* values are from 1-way ANOVA using the Dunnett adjustment for multiple comparisons. (**D**) Images of immunostaining for filaggrin (FLG, green) in cross-sections of control and KO organotypic cultures; bar = 50 µm; dashed lines mark bottom of the epidermis.

Immunoblotting lysates from KO epidermal cultures confirmed TEX264 depletion and showed no major compensatory changes in other reticulophagy receptors (**Figure 6B**). H&E staining revealed that KO organotypic epidermis lacked keratohyalin granules, similar to CX- and BafA1-treated cultures. Instead, the uppermost layers of KO cultures were filled with vacuoles, which can occur with blockade of autophagic degradation (16). Consistent with our data showing TEX264 induces nuclear shrinkage (**Figure 4E-F**), KO cultures lacking this reticulophagy receptor failed to undergo normal nuclear degradation and cornification (**Figure 6A, insets**).

Given the localization of endogenous TEX264 to the granular layers of the epidermis, we next investigated how loss of this reticulophagy receptor impacted filaggrin (FLG), a key component of the basophilic keratohyalin granules that heralds the final stage of keratinocyte maturation prior to cornification (48). FLG is a keratin-binding protein that aids in collapsing the intermediate filament cytoskeleton to allow flattening of cells as they transition into the cornified layers of the epidermis (45–47). Consistent with the visible depletion of keratohyalin granules in KO organotypic epidermis, cultures lacking TEX264 exhibited markedly reduced levels of FLG compared to controls (**Figure 6C-D**). Together, these results confirm that TEX264 plays an essential function in keratinocyte maturation and epidermal cornification.

### TEX264 is induced by SERCA2 inhibition and is dysregulated in Darier disease

To substantiate a role of TEX264 *in vivo*, we aimed to determine if this receptor contributes to a rare genetic skin cornification disorder with a known link to ER dysfunction. Darier disease (DD) is caused by dominant heterozygous mutations that deplete the sarco/endoplasmic reticulum calcium ATPase 2 (SERCA2) (22, 23), an ER calcium pump with a known role in autophagy (49–51). In fact, selective SERCA2 inhibitors, such as thapsigargin (TG), deplete ER calcium and induce protein misfolding, triggering autophagy and ER stress response pathways (52, 53). Previous work has shown that inducing ER stress in keratinocytes using TG or tunicamycin promotes keratinocyte differentiation (42, 54).

Treating organotypic cultures with TG to inhibit SERCA2 replicates features of DD (55), including precocious cornification, a diagnostic feature of DD pathology termed “dyskeratosis.” In organotypic epidermis, we found that TG accelerated the formation of granular cells and caused them to prematurely cornify, which decreased the average thickness of the epidermal tissue (**Figure 7A-B**), similar to our results from using tunicamycin to induce ER stress (**Figure 3**). Importantly, we linked TG-induced epidermal dysmaturation to earlier onset of TEX264, which was robustly induced in the intermediate cell layers of TG-treated cultures while controls restricted TEX264 to the granular layers (**Figure 7C-D**).

**Figure 7:**
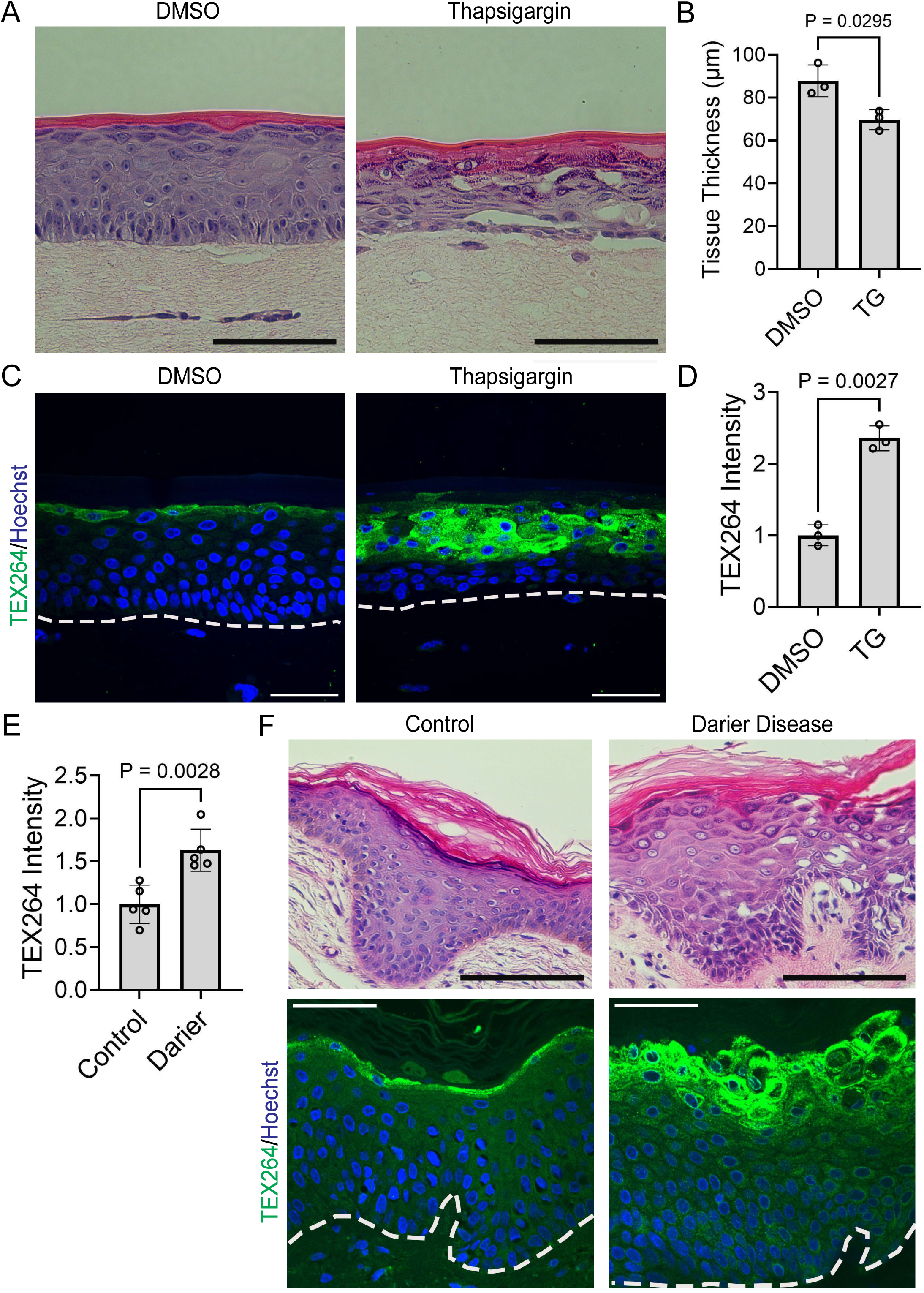
TEX264 is induced by SERCA2 inhibition and is dysregulated in Darier disease. **(A)** H&E-stained cross-sections of organotypic epidermal cultures treated with DMSO or thapsigargin (TG) for 48 hr; bar = 100 µm. (**B**) Bar graph displays the mean ± SD of the tissue thickness (µm) of epidermal cultures treated with DMSO or TG with individual data points representing the average thickness across N=3 biological replicates; *P* values from a ratio paired 2-tailed Student’s t test. (**C**) Immunostaining of TEX264 (green) and nuclei (Hoechst, blue) in cross-sections of organotypic epidermal cultures treated with DMSO or TG for 48 hr; bar = 50 µm; dashed lines mark bottom of epidermis. (**D**) Bar graph displays the mean ± SD of TEX264 fluorescence intensity in cultures treated with DMSO or TG with data points from N=3 biological replicates; mean intensity for DMSO normalized to 1; *P* value from a paired 2-tailed Student’s t test. (**E**) Bar graph displays the mean ± SD of TEX264 fluorescence intensity in biopsies from healthy control epidermis or lesional epidermis from Darier disease patients with data points from N=5 unrelated donors in each group; mean intensity for controls normalized to 1; *P* value from an unpaired 2-tailed Student’s t test. (**F**) H&E-stained cross-sections of biopsies from healthy control epidermis or lesional epidermis from Darier disease patients; bar = 100 µm; (below) immunostaining of TEX264 (green) and nuclei (Hoechst, blue) in cross-sections of biopsies from healthy control epidermis or lesional epidermis from Darier disease patients; bar = 50 µm; dashed lines mark bottom of the epidermis.

In DD, a reduced ability of SERCA2 to pump calcium ions into the ER is thought to lead to protein aggregation and ER stress (56–59), which can induce excessive autophagy and cell death (41, 60, 61). Consistent with a model in which SERCA2 loss-of-function triggers autophagy and premature cornification, we found in our cohort of de-identified biopsies from this rare disorder that lesions of DD exhibited an increase in TEX264 protein levels compared to normal control skin (**Figure 7E-F**). Indeed, whereas control epidermis restricted TEX264 to the granular layers that are preparing to cornify, in DD biopsies we noted TEX264 was more broadly expressed, particularly in areas of dyskeratosis, consistent with our finding that TEX264 induces premature cornification in cultured keratinocytes (**Figure 4**). Together, these data offer insight into the pathophysiology of DD: SERCA2 deficiency leads to ER stress, which induces premature expression of the reticulophagy receptor TEX264 and precocious cornification.

## DISCUSSION

To replenish the outermost protective skin layers, keratinocytes undergo progressive remodeling and degradation of their nuclei and organelles in a unique metamorphosis as they traverse the upper epidermis, flatten, and become corneocytes (3–5). Defects in cornification disrupt both the form and function of the skin in a group of diseases recently renamed epidermal differentiation disorders (EDDs) (62), but were originally termed ichthyosis due to accumulation of thick scales (8–10, 63). Many EDDs exhibit extra-cutaneous pathology, indicating that drivers of cornification can play crucial roles in other tissues (64, 65). While elucidating the biology of cornification is essential to design rational therapies for these orphan disorders, nearly all of which lack any FDA-approved therapy, understanding rare diseases can also reveal pathogenic mechanisms of more prevalent diseases (66). For example, prior work genetically linked impairment in autophagy to an ultra-rare EDD (67) and to the common skin disease, psoriasis (68–71), both characterized by thickened, scaly epidermis.

Autophagy is a catabolic pathway conserved from yeast to humans that can recycle cytoplasmic contents during starvation (72, 73) or eliminate damaged organelles accumulated during aging, particularly in neurodegeneration (74–76). This lysosomal degradation pathway is also responsible for programmed elimination of *normal* organelles during the differentiation and morphogenesis of specialized cells like erythrocytes (77, 78), Schwann cells (79), and ocular lens cells (80). Organelle breakdown and lysosomal engulfment were documented in terminally cornifying keratinocytes decades ago (12–14). Yet, the mechanisms controlling this process remain largely unknown, which has precluded targeted therapeutics to augment skin barrier function. Autophagy has emerged as an important regulator of the terminal phase of keratinocyte differentiation (70, 81–87); however, only recently has *selective* autophagy, driven by organelle-specific receptors (34, 35), been shown to participate in the wholesale clearing of keratinocyte nuclei and organelles (16, 88). While these studies demonstrated that mitochondrial elimination in the epidermis occurs via mitophagy, reticulophagy had not previously been known to operate in the skin.

Here, we show that the selective autophagy machinery is activated to break down the ER network at a critical point in the keratinocyte lifecycle just prior to cornification. Only about a decade ago did investigators identify a family of ER-localized autophagy receptors (26, 27) with TEX264 even more recently shown to be a *bona fide* reticulophagy receptor (37, 38). Interestingly, its full name, testis-expressed gene 264, derives from high expression of TEX264 in spermatocytes, which, similar to erythrocytes and keratinocytes, undergo selective organelle elimination as they mature into spermatids (89–91). We identified a critical function for TEX264 in keratinocytes, operating in the granular layers to trigger ER breakdown during skin barrier formation. Prior research showed autophagy can also degrade nuclear lamins (27, 92, 93), including in the epidermis (70, 94). As an ER-embedded protein, TEX264 can localize to the outer nuclear membrane and in HeLa cells, perinuclear TEX264 facilitated autophagic degradation of damaged DNA to support genome stability (43, 44). Intriguingly, we found TEX264 concentrated at the nuclear perimeter and it reduced nuclear size when ectopically expressed in differentiated keratinocytes. Whether this receptor plays a direct role in nuclear breakdown (i.e., nucleophagy) during cornification remains an area of active investigation.

Elegant work, including through intravital imaging of murine skin (19, 45, 95–97), has shown that the final stages of epidermal maturation are highly regulated to ensure homeostatic replacement of shed corneocytes to maintain epidermal thickness and preserve the skin barrier. We found that epidermal expression of TEX264 is tightly controlled, peaking in the granular keratinocyte layers poised to cornify. Our data show precocious TEX264 expression triggers premature ER breakdown and keratinocyte differentiation. As an added level of regulation, TEX264 contains serine residues upstream of its LIR domain that are phosphorylated by CK2 to enhance LC3 interaction (39). Treating TEX264-expressing keratinocytes with a CK2 inhibitor blocked ER fragment breakdown and impaired granular layer maturation. Moreover, TEX264-deficient cultures exhibited depletion of keratohyalin granules and filaggrin, a keratin-binding protein that facilitates cytoskeletal collapse during cornification (45–47, 96), substantiating a proposed specific role for autophagy in the granular layers (81, 84, 98).

We postulated that ER stress and the unfolded protein response (UPR), known inducers of reticulophagy (40, 99), may be upstream drivers of TEX264. Consistent with this, ER stress increases in the upper epidermal layers (17, 41, 42, 54, 100, 101), where TEX264 expression peaks. We showed that augmenting ER stress in organotypic epidermis using tunicamycin or thapsigargin (50, 102) caused precocious TEX264 expression. Both treatments accelerated the transition of keratinocytes into granular cells, causing premature cornification and epidermal thinning. Interestingly, thapsigargin blocks the ER calcium pump SERCA2, which is mutated in Darier disease (DD), a dermatologic disorder featuring premature cornification termed “dyskeratosis” (21–23). SERCA2 regulates autophagy in various cell types (50–52, 103), including keratinocytes, in which SERCA2 inhibition or knockdown impairs protein folding, inducing ER stress and the UPR (58, 59). Accordingly, excess ER stress and UPR activation have been linked to DD (55, 57, 104) as well as pemphigus, an autoimmune blistering disease with overlapping pathologic features (105). Keratinocytes from DD patients were recently found to have a reduced capacity for dampening reactive oxygen species, an output of sustained ER stress (56, 61, 99, 106). When the UPR is overwhelmed, cells can resort to apoptosis, which may manifest as dyskeratosis in DD lesions (60, 106).

In sum, our work supports a model of epidermal morphogenesis in which keratinocytes utilize selective autophagy receptors as cues to orchestrate the proper timing of organelle and nuclear breakdown to build the skin barrier. We defined a specific role for reticulophagy in keratinocyte differentiation through TEX264, an ER-localized selective autophagy receptor; its upregulation in the uppermost viable keratinocyte layers is an essential trigger for their terminal stage of maturation. We further demonstrated that SERCA2 loss-of-function augments TEX264, including in biopsies of DD, which may explain the aberrant keratinocyte differentiation that is a diagnostic feature of its pathology. Whether autophagy can be targeted for therapeutic benefit in disorders of cornification like DD remains to be tested. However, FDA-approved inducers of this pathway, including rapamycin, have been successfully deployed for off-label treatment of other skin diseases, including EDDs (68, 107–110).

## METHODS

### Sex as a biological variable

Our *in vitro* studies utilized available keratinocytes (primary and immortalized) from human males, but our results were validated in skin biopsy specimens from both male and female patients, which exhibited comparable findings.

### Reagents

SERCA2 inhibitor thapsigargin (Cat. #12758; 1 μM) was from Cell Signaling Technology. Casein kinase 2 inhibitor CX-4945 (Cat. #S2248; 10 μM) was from SelleckChem. Bafilomycin A1 (Cat. #B1793; 100 μM) and tunicamycin (Cat. #T7765; 5 μg/ml) were from Sigma. Dimethyl sulfoxide (DMSO; Cat. #BP231-100) was from Fisher Scientific.

Mouse anti-β-Actin (C4; Cat. #sc-47778; immunoblotting 1:500) and anti-GAPDH (0411; Cat. #sc-47724; immunoblotting 1:500) were from Santa Cruz. Mouse anti-RTN3/HAP was from Abcam (Cat. #ab68328; immunoblotting 1:200). Rabbit anti-TEX264 was from Sigma (Cat. #HPA017739; immunostaining 1:500) or ProteinTech (Cat. #25858-1-AP; immunoblotting 1:2000). Rabbit anti-filaggrin (Cat. #HPA030188; immunostaining 1:1000) and anti-SEC62 (Cat. #HPA014059; immunoblotting 1:500) were from Sigma. Rabbit anti-CCPG1 (E3C5G; Cat. #80158; immunoblotting 1:400) and anti-FAM134B (Cat. #61011; immunoblotting 1:1000) were from Cell Signaling. Chicken anti-calreticulin was from Abcam (Cat. #ab2908; immunostaining 1:1000). Hoechst 33342 was from Thermo-Fisher (Cat. #H1399; immunostaining 20 μg/ml).

Secondary antibodies for immunoblotting (1:10,000) were IRDye 800CW goat anti-rabbit IgG (Cat. #926-32211) and IRDye 680RD goat anti-mouse IgG (Cat. #926-68070) from LI-COR Biosciences. Secondary antibodies for immunostaining of cells and tissues (1:300) were from Thermo-Fisher: Goat anti-mouse IgG AlexaFluor-405 (Cat. #A31553), AlexaFluor-488 (Cat. #A11001), AlexaFluor-594 (Cat. #A11005), or AlexaFluor-633 (Cat. #A21050); goat anti-rabbit IgG AlexaFluor-405 (Cat. #A31556), AlexaFluor-488 (Cat. #A11008), AlexaFluor-594 (Cat. #A11012), or AlexaFluor-633 (Cat. #A21070); goat anti-chicken IgY AlexaFluor-488 (Cat. #A-11039), AlexaFluor-594 (Cat. #A-11042), or AlexaFluor-633 (Cat. #A-21103).

### Cell culture

All cell lines were cultured at 37°C in an air-jacketed humidified incubator with 5% CO2. Cells were cultured on sterile tissue culture-treated plates and passaged to remain sub-confluent using 0.25% Trypsin-EDTA (Thermo-Fisher, Cat. #15400054).

Normal human epidermal keratinocytes (NHEKs) were isolated from de-identified male neonatal foreskins by the Penn Skin Biology and Disease Resource-based Center. Cells were cultured in Medium 154 with 0.07 mM CaCl2 (Thermo-Fisher, Cat. #M154CF500) with 1x human keratinocyte growth supplement (Thermo-Fisher, Cat. #S0015) plus 1x gentamicin/amphotericin (Thermo-Fisher, Cat. #R01510).

hTERT-immortalized human epidermal keratinocytes (THEKs) from the original N/TERT-2G line (111) were cultured in keratinocyte serum-free medium (KSFM) from Thermo-Fisher (Cat. #37010022) with 0.2 ng/mL human epidermal growth factor, 30 µg/mL bovine pituitary extract plus 0.31 mM CaCl2, 100 U/mL penicillin, and 100 μg/mL streptomycin.

J2-3T3 immortalized murine fibroblasts (a gift from Dr. Kathleen Green, Northwestern University, Chicago, IL, USA) were grown in complete Dulbecco’s Modified Eagle Medium (DMEM) (Thermo-Fisher, Cat. #11965092) with 10% Hyclone FBS (Fisher Scientific, Cat. #SH3039603), 2 mM GlutaMAX (Thermo-Fisher, Cat. #35050061), plus 100 U/mL penicillin and 100 μg/mL streptomycin.

### CRISPR/Cas9 gene editing

CRISPR knockout (KO) keratinocytes were generated as described (112). Single-guide RNAs (sgRNAs) were designed to target *TEX264* (sgRNA: CATGTCGGACCTGCTACTAC) or the *TUBAP* pseudogene (sgRNA: GTATTCCGTGGGTGAACGGG) to generate a control KO line.

A web tool was used for CRISPR strategy (https://portals.broadinstitute.org/gpp/public/analysis-tools/sgrna-design). Synthetic sgRNA target sequences were inserted into a cloning backbone, pSpCas9 (BB)-2A-GFP (PX458) (Addgene, Cat. #48138), then were cloned into competent *E. coli* (Thermo-Fisher, Cat. #C737303). Proper insertion was validated using Sanger sequencing. Final plasmids were transfected into N/TERT-2G keratinocytes (111) using a TransfeX kit (ATCC, Cat. #ACS4005) with or without the JAK1/JAK2 inhibitor baricitinib (10 µg/mL). Single GFP-positive cells were plated then expanded. Sanger sequencing confirmed the presence of heterozygous or homozygous mutation in the targeted gene.

### Viral transduction

Human embryonic kidney (HEK) 293T retroviral packaging cells (Phoenix cells (113); a gift from Dr. Garry Nolan Lab, Stanford University, Palo Alto, CA, USA) or 293FT lentiviral packaging cells (a gift from Dr. Erika Holzbaur, University of Pennsylvania, Philadelphia, PA, USA) were cultured in complete DMEM. Transfections for retrovirus production used 4 μg retroviral plasmid DNA and for lentivirus used 2 μg lentiviral plasmid DNA plus 1 μg psPax2 (Addgene, Cat. #12260) plus 1 μg pMD2.G (VSV-G; Addgene, Cat. #12259). Plasmid DNA was incubated with 12 μL FuGENE 6 (Promega, Cat. #E2691) in 800 μL Opti-MEM (Thermo-Fisher Cat. #31985070) for 10 min at room temperature. The transfection cocktail was added dropwise onto sub-confluent Phoenix or 293FT cells in 60-mm dishes followed by incubation at 37°C.

After 24 hr, viral supernatants were collected and centrifuged at 200 g for 5 min to pellet any dislodged cells, then polybrene (Sigma, Cat. #H9268) was added (4 μg/mL). The supernatant was aliquoted and snap-frozen in liquid nitrogen for long-term storage at -80°C or used immediately. For transduction, native medium was removed from keratinocytes and replaced with viral medium for 1 hr at 37°C. After removing the viral medium and rinsing in PBS, native medium was replaced and cells were expanded in culture.

Retroviral constructs used for live fluorescent imaging (all gifts from Dr. Erika Holzbaur, University of Pennsylvania, Philadelphia, PA, USA) were made by sub-cloning the following plasmid inserts into the pLZRS retroviral vector: ER-dsRed2; ER-GFP; LC3B-mCherry; LAMP2-GFP; ER-GFP-RFP (Addgene, Cat. #128257). The TEX264 (Addgene, Cat. #128258) and TEX264-LIR (Addgene, Cat. #128259) inserts were sub-cloned into pLZRS with addition of a C-terminal EGFP. THEKs for imaging ER in organotypic epidermis (gifts from Drs. Andrew Kowalczyk and Navaneetha Bharathan, Penn State University, Hershey, PA, USA) were transduced with the pLenti6 lentiviral vector containing ER-StayGold (Addgene, Cat. #185822) or mChilada-VAPB (a gift from Nathan Shaner and Gerard Lambert, University of California San Diego School of Medicine, La Jolla, CA, USA).

### Organotypic epidermal cultures

Organotypic human epidermal “raft cultures” were generated as previously published (30, 114). Cultures grown from NHEKs were differentiated in E-medium, a 3:1 mixture of DMEM:Ham’s F12 (Thermo-Fisher, Cat. #11765054) supplemented with 10% FBS, 180 µM adenine (Sigma Cat. #A2786), 0.4 µg/mL hydrocortisone (Sigma, Cat. #H0888), 5 µg/mL human insulin (Sigma Cat. #91077C), 0.1 nM cholera toxin (Sigma, Cat. #C8052), 5 µg/mL apo-transferrin (Sigma Cat. #T1147), and 1.36 ng/mL 3,3′,5-tri-iodo-L-thyronine (Sigma Cat. #T6397). Cultures grown from THEKs were differentiated in CellnTEC Prime Epithelial 3D Medium (Zen-Bio, Cat. #CnT-PR-3D).

Dermal collagen rafts were made from J2-3T3 murine fibroblasts in transwells (Corning Cat. #353091). For each raft, 1 x 10^6^ fibroblasts were counted (Thermo Countess3, Cat. #AMQAX2000) and resuspended to 10% of the desired final volume using sterile reconstitution buffer (1.1 g of NaHCO3 and 2.39 g of HEPES in 50 mL 0.05 N NaOH). An additional 10% of the final desired volume of 10x DMEM (Sigma-Aldrich Cat. #D2429) was added. High-concentration rat tail collagen I (Corning, Cat. #CB354249) was then added (final concentration 4 mg/mL) and sterile water was used to dilute to the final volume, 2 mL total per raft. The collagen/fibroblast slurry was inverted to mix, then aliquoted as 2 mL per transwell insert placed within a deep 6-well culture plate (Corning, Cat. #08-774-183). The rafts were polymerized for 1 hr at 37°C, after which they were submerged in complete DMEM to incubate overnight in the 37°C incubator.

After 24 hr, DMEM was aspirated from both upper and lower transwell chambers and 1.5 x 10^6^ keratinocytes were seeded onto each raft in 2 mL of DMEM; additional DMEM was added to the bottom chamber to keep the raft submerged in liquid. Cultures were then incubated overnight at 37°C. The next day, DMEM was carefully removed from both transwell chambers and keratinocytes were placed at an air-liquid interface by adding either CnT 3D medium (for THEKs) or E-medium (for NHEKs) only into the lower chamber until reaching the bottom of the raft. Vehicle control (DMSO) or chemical inhibitors were diluted in the lower chamber medium as described in the figure legends. Organotypic cultures were matured for 5-12 days, replacing medium in the lower chamber every 2-3 days followed by live imaging or fixation. For histology, the transwell was transferred into a 6-well culture plate and submerged in 10% neutral-buffered formalin (Fisher Scientific, Cat. #22-026-435) for at least 24 hr.

### Tissue processing and histology

Organotypic epidermis was processed for histologic examination by Core A of the Penn SBDRC or the Experimental Histopathology Core of the Fred Hutchinson Cancer Center. Formalin-fixed paraffin-embedded cross-sections from human organotypic epidermal tissue or skin biopsies underwent standard histologic processing and hematoxylin and eosin (H&E) staining. A 40x long working-distance, achromatic, phase-contrast objective on the EVOS FL microscope (Thermo-Fisher) was used to obtain H&E images with the high-sensitivity embedded interline CCD color camera.

### Tissue thickness quantification

Thickness measurement of epidermal tissues was performed by hand using non-visibly labeled H&E images. The “Polygon” tool in Fiji (ImageJ) was used to encircle the entire epidermis in multiple non-overlapping fields across multiple tissue sections for each condition. The “Measure” function was used to calculate the area of the encircled epidermis, which was divided by its length to calculate the average thickness across multiple independent experiments.

### Immunoblotting

Whole-cell lysates of keratinocytes or organotypic cultures were made using urea sample buffer (USB) [8 M Urea, 1% SDS, 10% glycerol, 0.0005% pyronin-Y, 5% β-mercaptoethanol, 60 mM Tris, pH 6.8] and were homogenized using a microtip probe sonicator (Fisher Scientific). Lysates were separated by electrophoresis at 80V for 1 hr in NuPAGE MES-SDS Running Buffer (Thermo-Fisher, Cat. #NP0002) and loaded into NuPAGE 12% Bis-Tris Gels (Thermo-Fisher, Cat. #NP0343BOX). Proteins were transferred for 60 min at 50 V onto Immobilon-FL membrane (Millipore Cat., #IPFL85R) using transfer buffer (25 mM Tris, 192 mM glycine, 20% (v/v) methanol). Membranes were dried overnight. The next day, blots were re-wetted in 100% methanol and re-hydrated in Tris-buffered saline (TBS). Revert total protein stain and Revert wash solution (LI-COR, Cat. #926-10016) were used for total protein staining captured on an Odyssey M Imaging System followed by incubation in Revert destaining solution and rinsing in water. Membranes were blocked for 60 min at room temperature in Intercept PBS blocking buffer (LI-COR, Cat. #927-70003), then were probed at 4°C overnight in primary antibodies in Intercept PBS blocking buffer with gentle rocking. Blots were washed three times in 1x TBS with 0.1% (v/v) Tween-20 (TBS-T), then were incubated 1 hr at room temperature with gentle rocking in dark boxes in Intercept TBS blocking buffer with 0.02% Tween-20 and 0.01% SDS plus fluorescent secondary antibodies (LI-COR) at 1:10,000. Blots were washed three times in TBS-T then scanned on the Odyssey M.

### Fluorescent immunostaining of cells

Keratinocytes were grown in 35-mm glass-bottom cell culture dishes (MatTek, Cat. #P35G-1.5-20-C) and fixed in 4% paraformaldehyde at 37°C for 10 min followed by rinsing in phosphate-buffered saline (PBS). Cells were permeabilized in 0.2% Triton in PBS at room temperature for 5 min. Fixed cells were incubated 30 min at 37°C in blocking solution [0.5% (w/v) bovine serum albumin (BSA, Sigma, Cat. #A9647), 10% (w/v) normal goat serum (NGS, Sigma, Cat. # G9023) in PBS]. Cells were rinsed with PBS then incubated in primary antibodies diluted in 0.5% (w/v) BSA in PBS overnight at 4°C. Cells were rinsed in PBS 3 times, then were incubated in secondary antibodies (+/- Hoechst) in 0.5% (w/v) BSA in PBS for 60 min at 37°C. Cells were rinsed three times in PBS and held in PBS for confocal microscopy.

### Fluorescent immunostaining of tissues

Formalin-fixed paraffin-embedded tissue cross-sections on glass slides were baked for at least 2 hr at 65°C. Slides were then immersed for 3 min in each of: 3 baths of xylenes (Fisher, Cat. #X3P), 3 baths of 95% ethanol, 1 bath of 70% ethanol, and 3 baths of PBS. Slides were submerged in antigen retrieval buffer [0.1 M sodium citrate (pH 6.0) with 0.05% (v/v) Tween-20] and heated to 95°C for 15 min. After cooling to room temperature, slides were rinsed once in PBS. A hydrophobic barrier pen was used to encircle tissue sections. Tissue sections were incubated in blocking buffer [0.5% (w/v) BSA, 10% (v/v) NGS in PBS] within a humidified chamber for 30 min at 37°C. Slides were rinsed for 3 min in each of 3 PBS baths, then were incubated overnight at 4°C in primary antibodies diluted in 0.5% (w/v) BSA in PBS in a humidified chamber. Slides were then washed in 3 baths of PBS for 3 min each then were incubated for 60 min at 37°C in secondary antibodies (+/- Hoechst) in 0.5% (w/v) BSA in PBS in a humidified chamber. Slides were washed in 3 baths of PBS for 3 min each. Finally, Prolong Gold (Thermo-Fisher, Cat. #P36934) was applied to cover the tissue sections under a #1.5 glass coverslip. After drying, slides were held in the dark prior to imaging on a confocal microscope.

### Confocal fluorescence microscopy

A Yokogawa W1 spinning-disk confocal (SDC) system on a Nikon Ti2 microscope with a Hamamatsu ORCA-FusionBT sCMOS camera was used to acquire images. Samples were illuminated using laser excitation (405, 488, 561, 640 nm) and emitted fluorescence was captured through a 40x 0.95 NA air objective or 60x 1.2 NA water objective (Nikon) and standard filters. For confocal imaging of live submerged cultures, cells transduced with fluorophore-tagged constructs were seeded into 35-mm glass-bottom dishes at least 24 hr prior to imaging in their native medium at 37°C in a stage-top environmental chamber with a sliding lid (Okolab). For imaging of live stratified epidermis, organotypic cultures were transferred from transwells into 35-mm glass-bottom dishes with the apical tissue surface on the glass. Z-stack images through the epidermal layers were acquired using a piezo motor step size of 200 nm.

### Fluorescent immunostaining quantification

Immunostained cell sheets or tissue sections were imaged using SDC microscopy as above. Fluorescence intensity was quantified using Fiji to display non-visibly labeled images of immunostained cells or tissues. The Fiji “Measure” function was used to calculate the mean fluorescence intensity across the entire visual field (for confluent cell sheets) or within a circumscribed region of interest (ROI) using the “Polygon” tool to encircle a single cell (for ER degradation assessment) or the entire epidermis (for tissue cross-sections of organotypic cultures or skin biopsies); the same field or ROI was used to measure the mean fluorescence intensity for each laser channel in multi-color images. The mean fluorescence intensity from each image was averaged for each condition and in multiple experimental replicates for statistical analysis; the average intensity of pooled control samples was normalized to be 1.

### Perinuclear enrichment assessment

Fiji was used to calculate perinuclear enrichment of GFP signal in 60x SDC images of keratinocytes. Paired 100 x 100 pixel ROIs within a single cell were selected from the cytoplasm versus across the nuclear boundary. The Fiji “Measure” function was used to calculate the mean fluorescence intensity within each ROI and a perinuclear enrichment factor was calculated as the ratio of the perinuclear-to-cytoplasmic mean intensity. This ratio was averaged across pooled cells from each condition in multiple independent experiments for statistical analysis.

### Nuclear size quantification

Nuclear cross-sectional area was measured using the Fiji MorphoJ plug-in on 60x SDC images from the uppermost layer of GFP-positive THEKs that were differentiated in CnT 3D medium for 4 days followed by paraformaldehyde fixation and staining with Hoechst. If needed, the “Adjustable Watershed” tool was used to segregate adjacent nuclei; overlapping nuclei, mitotic cells, and cells with multiple nuclei were excluded from the analysis. MorphoJ outlined each nucleus and the BIOP plugin assigned a unique ID to each nucleus. The mean nuclear cross-sectional area was calculated using the MorphoJ “Analyze” function and was averaged across multiple independent experiments for each condition.

### Statistics

Statistical analyses and data graphing were performed using Prism version 9 (GraphPad). Statistical parameters including sample size, center definition, measures of dispersion, and statistical tests are included in each figure legend. Means of 2 normally distributed groups were compared with a two-tailed unpaired or paired Student’s t test. Means from ≥2 normally distributed groups were compared using one-way ANOVA with *P* values adjusted for multiple comparisons. *P* values <0.05 were considered statistically significant. Graphs include exact *P* values.

### Study approval

Normal human epidermal keratinocytes (NHEKs) from neonatal foreskins and human skin biopsies were procured by the Penn Skin Biology and Diseases Resource-based Center (SBDRC) in a de-identified manner under protocols (#808224; #808225) approved by the Institutional Review Board (IRB) of the University of Pennsylvania. Use of de-identified cells and tissues (collected for clinical purposes that would otherwise be discarded) was exempted by the IRB for written informed consent.

### Data availability

All underlying values for graphed data are available in the **Supporting Data Values** file.

## AUTHOR CONTRIBUTIONS

Conceptualization: C.J.J., A.T., C.L.S. Data Curation: C.J.J, C.L.S. Formal analysis: C.J.J., E.A., R.K., C.L.S. Investigation: C.J.J., A.T., A.S.P., B.D.L., A.R., A.C., M.K.S., C.L.S. Resources: J.E.G., C.L.S. Supervision: J.E.G., C.L.S. Visualization: C.J.J, A.T., A.S.P., B.D.L., A.R., C.L.S. Validation: A.T., A.C, M.K.S., C.L.S. Writing (original draft): C.J.J. Writing (review and editing): C.J.J., C.L.S.

## ACKNOWLEDGEMENTS

We thank the Skin Biology and Diseases Resource-Based Centers at University of Pennsylvania (NIH P30-AR069589) and University of Michigan (NIH P30-AR075043) for keratinocytes, CRISPR/Cas9 editing, histology, and archived biopsies sections. C.L.S. acknowledges grant support from NIH K08-AR075846, NIH R03-TR005428, the American Skin Association, and the LEO Foundation (LF-OC-23-001393). C.J.J. was supported by the American Society for Clinical Investigation Physician-Scientist Support Foundation Fellowship. A.T. was supported by NIH R03-AR082896 and an Innovation Pilot Award from the University of Washington Institute for Stem Cell and Regenerative Medicine. A.P. was supported by a grant from the Foundation for Ichthyosis and Related Skin Types (FIRST). J.E.G., M.K.S., and A.C. were supported by NIH P30-AR075043, the Taubman Medical Research Institute, and the National Psoriasis Foundation (852098). BioRender.com was used to make the **Graphical Abstract** and **Figures 1A, 1C, and 2E**. We thank Mariko Tokito and Erika Holzbaur (University of Pennsylvania, Philadelphia, PA, USA) and Andrew Kowalczyk and Navaneetha Bharathan (Penn State University, Hershey, PA, USA) for reagents and experimental advice.

